# A phylogenomic and metagenomic meta-analysis of bacterial diversity in the phyllosphere lifts a veil on Hyphomicrobiales dark matter

**DOI:** 10.1101/2025.08.19.671110

**Authors:** Jean-Baptiste Leducq, Louis-Philippe St-Amand, David Ross, Steven W. Kembel

## Abstract

The phyllosphere, or above-ground part of plants, hosts diverse bacterial communities that play critical ecological roles and provide beneficial functions for the plant. The Hyphomicrobiales (Alphaproteobacteria) are a highly diverse and ecologically important clade known to be key members of the plant microbiome, in particular in association with plant roots, but their diversity remain largely uncharacterized in the phyllosphere. Using a meta-analysis combining metabarcoding, metagenomics and phylogenomics, we explored the worldwide diversity of leaf-associated Hyphomicrobiales. We confirmed *Methylobacterium* was ubiquitous in the phyllosphere and revealed the dominance of two under-characterized Hyphomicrobiales taxa: *Lichenibacterium*, a lichen-associated genus previously identified as “1174-901-12” in taxonomic databases, and RH-AL1, an undescribed lineage of bacteria related to Beijerinckiaceae, previously isolated from coal slag. Despite their abundance in the phyllosphere, *Lichenibacterium* and RH_AL1 could not be properly detected by 16S rRNA gene barcoding, due in part to limitations of taxonomic resolution of the 16S rRNA gene and of representativeness in existing taxonomic databases, underlining limitations of this approach for their accurate identification in the phyllosphere. As for *Methylobacterium,* a significant proportion of *Lichenibacterium* and RH-AL1 were also detected in association with lichens and in environments with harsh conditions like exposed surfaces, air and snow, suggesting airborne or waterborne dispersal and high resilience in harsh environments. Overall, our study stresses the need to move toward metagenomics and culturomics to increase the representativeness of leaf-associated bacterial taxa in reference databases, and to improve our understanding of the evolutionary and functional mechanisms underpinning bacteria adaptations to living on plants.

**Graphical abstract:** 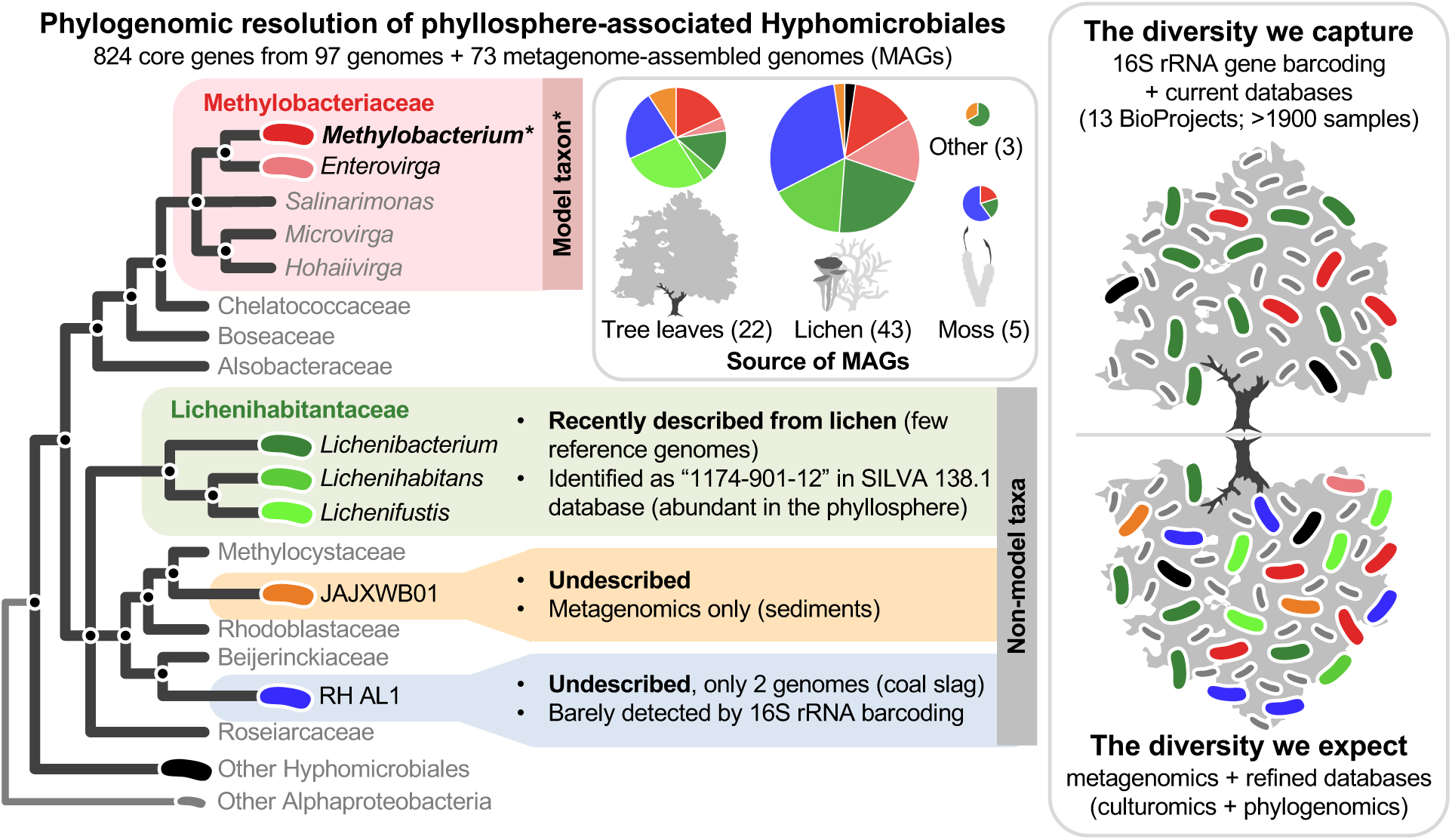

**Highlights:**

- Global meta-analysis reveals core Hyphomicrobiales in the phyllosphere
- *Lichenibacterium* and *Methylobacterium* dominate the phyllosphere and exposed surfaces
- Novel phyllosphere clades RH-AL1 and JAJXWB01 were identified by phylogenomics
- 16S rRNA gene limits taxonomic resolution in the phyllosphere; metagenomics refines taxonomy

## INTRODUCTION

The phyllosphere, or the aerial parts of plants including leaves, has been intensively studied due to its crucial role in global carbon and nitrogen cycles. Recent advances in microbial ecology also revealed that, despite exposure to UV radiation, drought, and rapid environmental fluctuations, plant leaves host a large microbial diversity providing essential functions for plant health including nutrient acquisition, deterring pathogens, and facilitating resilience to stresses [41]. The phyllosphere microbiota can also modulate leaf surface chemical composition by releasing plant growth promoters such as vitamins and hormones, and by degrading toxic compounds such as pesticides [48]. Our knowledge about beneficial plant-associated bacterial diversity has been dominated by studies of the rhizosphere, and the phyllosphere was less well studied until recently. Roots are less exposed to environmental variation and are more nutrient-rich habitats than leaves. Therefore, root-associated microbes appear to be more rich and abundant, and microbial diversity and culture conditions are more predictable in the rhizosphere than in the phyllosphere [12]. For instance, Hyphomicrobiales (previously Rhizobiales) is a major plant-associated bacterial order that includes the intensively studied nitrogen-fixing bacteria in root nodules [13,43], while the phyllosphere-associated Hyphomicrobiales are mostly known for *Methylobacterium* (Methylobacteriaceae), a ubiquitous group of bacteria metabolizing one-carbon compounds such as methanol that are released by plants during cell-wall synthesis [39]. As a result, only a tiny fraction of leaf-associated microbes have been studied and cultivated so far, stressing the need for more studies that consider the phyllosphere, and for culture conditions that maximize their isolation success, such as those developed for the *Arabidopsis* microbiome [6]. Since the advent of molecular biology, microbial taxonomy has been based on universal marker genes (i.e. the 16S rRNA gene in bacteria), allowing identification of major taxonomic groups and their phylogenetic relationships [29]. Unfortunately, those markers do not give the resolution required to confidently define lower taxonomic units (genus, species), introducing unwanted artefacts in the phylogeny and classification of major groups of plant-associated microbes including *Methylobacterium* [2]. Alternatively, the *rpoB* gene has higher resolution [28] and has been used as a molecular marker to infer *Methylobacterium* diversity in the phyllosphere, with potential use at a larger taxonomic scale in the Hyphomicrobiales [20]. Overall, metabarcoding has allowed the quantification of microbial diversity without using cultivation but is still limited by using a single marker gene. Alternatively, shotgun metagenomics is a promising approach to infer the taxonomy, function and diversity of phyllosphere bacterial communities. This approach is however limited by the overall low bacterial abundance on the surface of plant leaves, and by the presence of high concentrations of plant host DNA for endophytic organisms. Finally, whole genome sequencing (WGS) of isolates, including plant-associated bacteria, coupled with phylogenomics (whole-genome-based phylogeny) substantially improved the taxonomy of Hyphomicrobiales in recent years [13,17]. Regardless of the method, taxonomic databases are still biased toward model taxa like *Methylobacterium*, and we argue that major phyllosphere-associated Hyphomicrobiales taxa are still omitted, underrepresented or misclassified in these databases. For instance, *1174-901-12* is an uncultured Hyphomicrobiales taxon identified in the 16S rRNA gene SILVA database (v138.1) [33] that has been consistently detected worldwide on the surface of plant leaves [3,16,19,20,23,36,44,46], where it typically represents 3 to 22% of total bacteria abundance according to amplicon sequencing data. Little is known about the taxonomy and the ecological role of *1174-901-12*, although its detection in lichens [35], in aerosols [25] and on various exposed surfaces [4,26,34,40] suggests a broader ecological spectrum than the phyllosphere. According to the SILVA database (v138.1) based upon the 16S rRNA gene, *1174-901-12* was member of the Beijerinckiaceae, a catch-all family of phyllosphere-associated Hyphomicrobiales also including *Methylocella*, *Methylocystis*, *Methylobacterium* and *Roseiarcus*, for instance (Table 1). Those genera were also classified in Beijerinckiaceae according to the Genome Taxonomy Database (GTDB; Table 1), while according to NCBI taxonomy database and to the List of Prokaryotic names with Standing in Nomenclature (LPSN), they belong to distinct families (Table 1). The underrepresentation and misclassification of major phyllosphere-associated Hyphomicrobiales such as *Methylobacterium* and *1174-901-12* illustrates the limits of using the 16S rRNA gene in describing the composition of microbial communities in the phyllosphere. It also stresses the need to move towards a metagenomic approach to enrich databases with genomes from uncultured taxa, and to a phylogenomic-based approach to clarify their taxonomy.

**Table 1:**
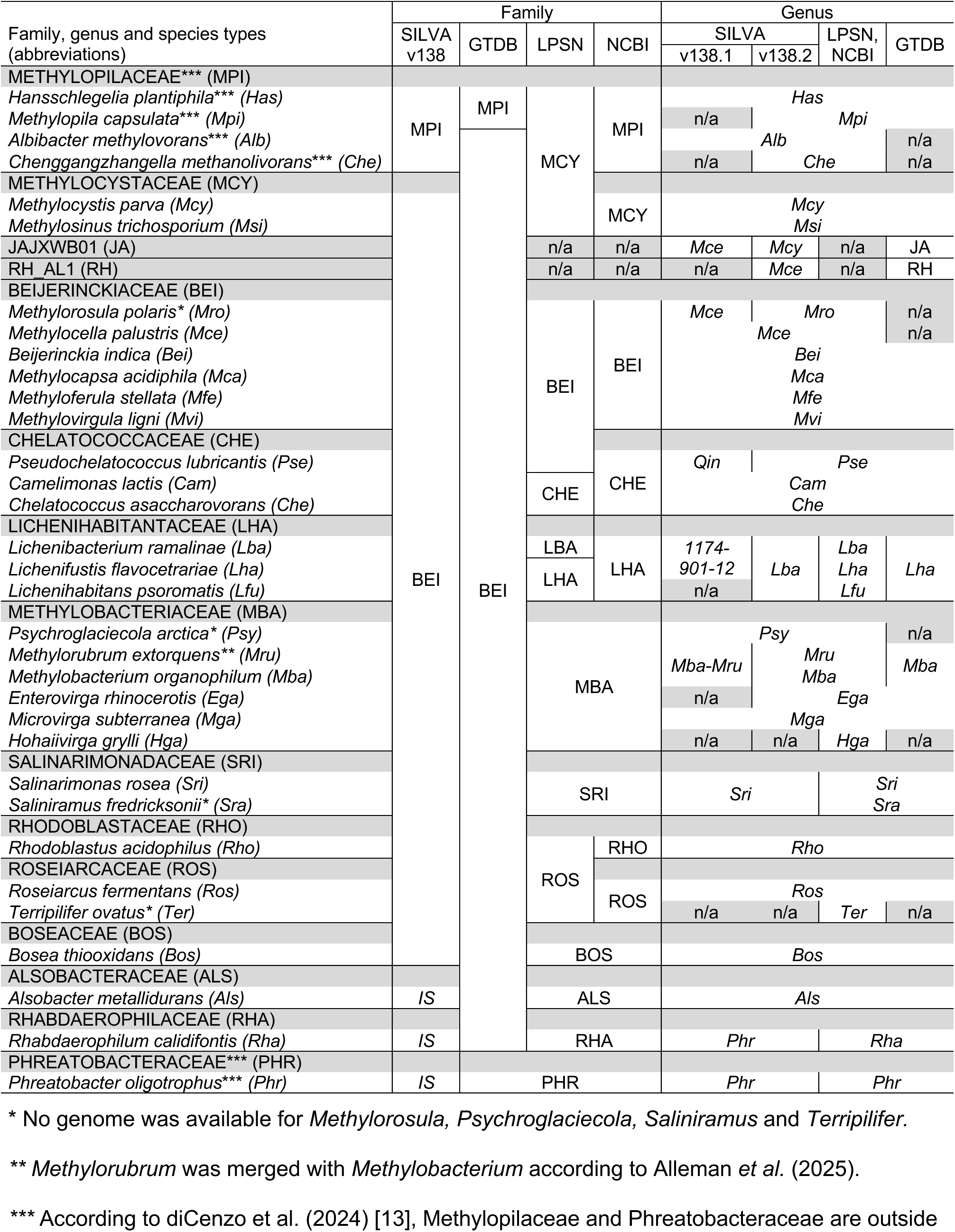
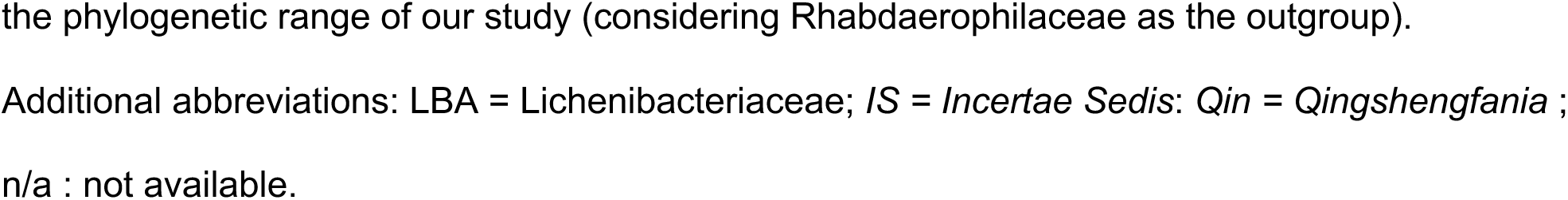
Comparison of taxonomic classifications (Family and Genus) of type strains from phyllosphere-associated Hyphomicrobiales (syn. Rhizobiales in GTDB and SILVA 138.1) with following databases: LPSN, NCBI, GTDB, and SILVA. Left columns: genus and species type names follow LPSN classification; family follow previous phylogenomic-based classification (diCenzo et al. 2024) [13]. Taxa not considered for phylogenomic analysis are indicated by asterisks (see footnotes for details).

Using a meta-analysis combining metabarcoding, metagenomics and phylogenomics, we explored worldwide the diversity of leaf-associated Hyphomicrobiales, incorporating data on microbial associations with vascular plants, mosses, and lichens. We confirmed the ubiquity of *Methylobacterium* in the phyllosphere and revealed the dominance of two under-characterized Hyphomicrobiales taxa: *Lichenibacterium*, a recently described genus of lichen-associated bacteria formerly identified as *1174-901-12*, and RH-AL1, an undescribed genus of bacteria related to Beijerinckiaceae, previously isolated from coal slag. The fact that *Lichenibacterium* and RH_AL1 were not properly detected by classical 16S rRNA gene barcoding despite their abundance in the phyllosphere illustrates the limitations of this metabarcoding approach, likely due to the lack of reference sequences for some taxa or they misclassification in reference databases, and the lack of taxonomic resolution for this marker gene. Like for *Methylobacterium,* a significant proportion of *Lichenibacterium* and RH-AL1 were also detected in lichen and in environments with harsh conditions like exposed surfaces, air and snow, suggesting airborne or waterborne dispersal and high resilience in adverse environments. Overall, our study stresses the biases of classical tools and reference database, as well as the power of metagenomics and phylogenomic to reveal uncultured leaf-associated bacteria. Future works should also use such tools, as well as culturomics, to increase the representation of these taxa in reference databases, and to address evolutionary and functional mechanisms underpinning bacterial adaptation to life on plants.

## METHODS

### Identification of 1174-901-12 as Lichenihabitantaceae

Because taxonomy of the phyllosphere-associated taxa *1174-901-12* identified in the SILVA database (v138.1) was unclear, we performed a preliminary phylogenetic analysis of 16S rRNA sequences used to define this taxon in SILVA v138.1 database, in order to validate that it corresponded to *Lichenibacterium*, as identified in the SILVA v138.2 database. *Lichenibacterium* [30] is part of Lichenihabitantaceae, a recently described lichen-associated family of bacteria, also including genera *Lichenifustis* [31], *and Lichenihabitans* [27]. For this analysis, we considered the 24 sequences identified as *1174-901-12* in the SILVA v138.1 database, 24 complete 16S rRNA gene sequences available for the family Lichenihabitantaceae in NCBI database, including reference strains, and two sequences from *Roseiarcus* as outgroup (Dataset 2). The Maximum-Likelihood tree was inferred with RAxML (1,000 bootstraps, GTRGAMMA).

### Comparison of phyllosphere-associated Hyphomicrobiales taxonomies among databases

We used the most recent phylogenomic-based survey of Hyphomicrobiales by diCenzo *et al.* (2024) [13] (syn. Rhizobiales) as a reference for taxonomy (Table 1; Dataset 1). According to this study, we limited the taxonomic range to a monophyletic group embedded within Hyphomicrobiales, and including genera *Methylobacterium* (family Methylobacteriaceae) and *Lichenibacterium*, AKA *1174-901-12* (Lichenihabitantaceae). We will refer to this group as phyllosphere-associated Hyphomicrobiales. This monophyletic group also included families Roseiarceae, Rhodoblastaceae, Methylocystaceae, Beijerinckiaceae, Chelatococcaceae, Salinarimonadaceae, Boseaceae and Alsobacteraceae (Table 1). Still according to diCenzo *et al.* (2024) [13], we used Rhabdaerophilaceae as the outgroup [13]. For each family cited above and embedded genera, we retrieved type species and strains from LPSN and compared they taxonomic classification (Family and Genus) between LPSN, NCBI, GTDB and SILVA databases (Table 1). In order to highlight incongruences between databases, we also included any other genera classified in above-mentioned families by at least one database, hence including genera from family Methylopilaceae, classified as Methylocystaceae in LPSN, and Phreatobacteraceae, to which Rhabdaerophilaceae was assigned with SILVA (Table 1). For SILVA, taxonomic assignment was performed on complete 16S rRNA sequences from type strains retrieved from LPSN, and from annotated genome assemblies for two phyllosphere-associated taxa exclusively reported in the GTDB database (RH_AL1 and JAJXWB01; see next section). Taxonomic assignment was performed with R package dada2 (function *assignTaxonomy* [10]) using SILVA v138.1 and v138.2 as references.

### Metagenome and genome collection of representative Hyphomicrobiales taxa from the phyllosphere, moss and lichen

We assembled a collection of phyllosphere-associated Hyphomicrobiales (syn. Rhizobiales) reference genomes (*n* = 97), and any metagenome-assembled genome (MAGs; n = 73) publicly available in NCBI databases. MAGs were mostly identified in three metagenomic surveys on microbial diversity at the surface of tree leaves, mosses, lichen, and their taxonomy was previously assessed with the GTDB database and assigned to Beijerinckiaceae (Rhizobiales). Leaf-associated MAGs (n=22) were identified in samples from the Canadian temperate forest (unpublished data; bioproject PRJEB93989; **Dataset 3**) and originally assigned to genera *Methylobacterium* (n= 4), *Enterovirga* (n=1), *Lichenihabitans* (n= 8), to undescribed genera RH-AL1 (n= 5) and JAJXWB01 (n= 2), or were unidentified at the genus level (n=2; **Dataset 3).** Lichen-associated MAGs came from a worldwide survey (n=43; **Dataset 3** [38]) and were previously assigned to genera *Lichenihabitans* (n=16), *Enterovirga* (n= 6), *Methylobacterium* (n= 6), RH_AL1 (n= 13) or were unidentified at the genus level (n=2; **Dataset 3)**. Moss-associated MAGs were previously identified in samples from the Canadian boreal forest (n=5; **Dataset 3** [18]) and were originally assigned to genera *Lichenihabitans* (n=1), *Methylobacterium* (n=1) and RH_AL1 (n=3). Two additional MAGs, previously assigned to genera *Lichenihabitans*, were identified in glacier sediments [9] and in mining soil [24].

We retrieved 97 reference genomes from NCBI, spanning the taxonomic and phylogenic range of Hyphomicrobiales-associated taxa (Table 1; **Dataset 3**). For Methylobacteriaceae, we maximized the genomic diversity of reference genomes by selecting the best assembly from each lineage previously identified in *Methylobacterium* (n=23; [2]), *Enterovirga* (n=3), *Microvirga* (n=2) and included the genome of a newly described genera, *Hohaiivirga* (n=1; [42]; **Dataset 3**). Because other leaf-associated taxa had fewer representatives in genomic databases, we retrieved all available genomes for Lichenihabitantaceae (n=8) and RH_AL1, only represented by two strains isolated from coal slag with genomes available in NCBI [45]. JAJXWB01 was previously identified by metagenomics in mine drainage sediments (unpublished data; bioproject PRJNA666025) and had no reference genome in NCBI. Accordingly, we included the JAJXWB01 assembly to our MAG collection (**Dataset 3**). To mitigate biases in phylogenomic reconstruction of the whole group, we also included NCBI RefSeq genomes from families that were not identified in leaf-associated MAGs, but were also members of this monophyletic group [13]: Methylocystaceae (n=12), Beijerinckiaceae (n=10), Roseiarcaceae (n=1), Rhodoblastaceae (n=3), Chelatococcaceae (n=11), Alsobacteraceae (n=3), Boseaceae (n=14), Salinarimonadaceae (n=3) and Rhabdaerophilaceae as the closest known outgroup with available genome assembly, (n=1; **Dataset 3**).

### Phylogenomic reconstruction of phyllosphere-associated Hyphomicrobiales

In order to clarify their phylogenetic relationships and taxonomy, we reconstructed the phylogenetic tree of phyllosphere-associated Hyphomicrobiales as described previously for Methylobacteriaceae [21] with the following modifications. Briefly, we annotated *de novo* the 97 genomes and 73 MAGs using RAST [5] and identified core genes (i.e. genes present in a single copy in all genomes), excluding hypothetical proteins. Because our analysis included highly fragmented MAGs, we used more relaxed criteria to define the core genome. First, we considered any gene present on average in 2 copies or less in at least 80% of genome assemblies as candidate core genes (MAGs excluded). Second, we estimated the real copy number of each candidate core gene in each assembly. As the proxy for the copy number, we normalized the total size of gene copies observed for a given gene in each assembly by the average size of all copies observed across all assemblies. We considered any normalized value lower than 0.7 (missing or too fragmented copy) or above 1.3 (likely duplication) as missing data. Third, we only considered candidate core genes with missing data in less than 30% of assemblies as core genes. We performed sequence alignment for each core gene individually, with the R package *msa* [8], using nucleotide sequence for rna-encoding genes and amino-acid sequence for protein-encoding genes. We computed a maximum-likelihood phylogenetic tree for each core gene independently using RAxML (GTRGAMMA model, 1,000 replicated trees [37]). We reconstructed the phyllosphere-associated Hyphomicrobiales consensus tree using ASTRAL-III, a coalescent-based method combining ML trees determined for each core gene independently, accounting for incomplete lineage sorting and horizontal gene transfers [47].

### Meta-analysis of barcoding-based phyllosphere studies

We selected previously published metabarcoding data based on the 16S rRNA gene from the tree phyllosphere and potentially related microbial environments and sources (other plants, mosses, lichen, exposed surfaces, air, snow). We restricted our survey to 13 recent studies for which BioProjects and related metadata were publicly available (**Dataset 4**). For each BioProject, we processed 16S rRNA gene reads to obtain an ASV (Amplicon Sequence Variants) abundance table per sample, using the package *dada2* in R [10] as described previously [20], with the following modifications. According to sequence quality profiles specific to each BioProject, 3’ ends of forward and reverse reads from each sequence were trimmed (option truncLen in function *filterAndTrim*) at specific lengths (**Dataset 4**). Additionally, 5’ ends of both reads were trimmed 20bp to remove the primer part (option trimLeft in function *filterAndTrim*). To correct for the great variation in read number among samples and BioProjects, we performed rarefaction curves to determine a conservative number of sequences to conserve per sample. Accordingly, we randomly selected 3,000 sequences per sample, hence excluding samples with fewer than 3,000 sequences (**Dataset 4**). We pooled together unique ASV sequences found among all BioProjects after rarefaction (N=57,100 unique sequences) and inferred their taxonomy in two steps. We first classified all ASVs to the genus level with the SILVA 138.2 database [33] using the *assignTaxonomy* function (R package dada2). To verify SILVA v 138.2 accuracy in predicting phyllosphere-associated Hyphomicrobiales ASVs, we extracted 16S rRNA gene sequences from complete phyllosphere-associated Hyphomicrobiales reference genomes (see section *Phylogenomic reconstruction of phyllosphere-associated Hyphomicrobiales*) and classified them as we did for ASVs. We also performed a phylogenetic analysis of 16S rRNA sequences extracted from reference genomes for a comparison with our phylogenomic reconstruction. The Maximum-Likelihood tree was inferred with RAxML (1,000 bootstraps, GTRGAMMA). For the second step, we refined ASV classification restricted to phyllosphere-associated Hyphomicrobiales sequences (i.e. identified as Beijerinckiaceae, *Alsobacter* or *Phreatobacter* with SILVA v138.2 database; Table 1) using the *assignTaxonomy* function with 16S rRNA gene sequences we extracted from complete phyllosphere-associated Hyphomicrobiales reference genomes as references. We subdivided Bioprojects from different climatic regions, or with samples from different substrates, in different categories. We subdivided Bioproject PRJNA100102 in three categories: Temperate forest, Subtropical forest and Tropical forest [44]. We subdivided Bioproject PRJNA290145 in three categories: Bark samples, Lichen samples and Moss Samples [4]. For each BioProject or category within BioProjects (n=17), we estimated the diversity of bacteria taxon as the relative abundance of sequences assigned to each taxon, focusing on Hyphomicrobiales dominant families and nested genera.

### Comparison of 16S rRNA and *rpoB*–based barcoding for the identification of phyllosphere-associated Hyphomicrobiales taxa

To assess the performance of the *rpoB* gene as a phylogenetic and taxonomic marker for phyllosphere-associated *Hyphomicrobiales*, we first retrieved complete *rpoB* gene sequences from reference genomes and MAGs with known taxonomy. *rpoB* sequences were extracted from RAST output files, aligned, and their phylogeny was assessed with RAxML as previously described for core genes. To compare *rpoB*- and 16S rRNA gene–based taxonomic resolution, we used amplicon data we previously generated [20], including 184 phyllosphere samples collected from two temperate forests in Quebec, Canada: *Station Biologique des Laurentides* (SBL, n = 99) and *Mont St-Hilaire* (MSH, n = 85). All samples were previously amplified with primers optimized for Methylobacteriaceae and targeting a hypervariable region of the *rpoB* gene [20]. We used *rpoB* ASV abundance tables and taxonomy from our previous study as a starting point. We reclassified ASVs using *rpoB* sequences from genomes and MAGs as references, excluding sequences that did not completely overlapped ASV sequences. Because reference sequences only included phyllosphere-associated Hyphomicrobiales, we only performed reclassification for ASVs previously identified as Rhizobiales (former Hyphomicrobiales), hence excluding Caulobacterales, Sphingomonadales and Rhodospirillales. Within Rhizobiales, we kept ASVs previously assigned to phyllosphere-associated Hyphomicrobiales (Methylobacteriaceae, Lichenibacteriaceae, Beijerinckiaceae, Methylocystaceae), hence excluding ASVs previously assigned to other families (Aurantimonadaceae, Bradyrhizobiaceae, Brucellaceae, Hyphomicrobiaceae, Phyllobacteriaceae and Rhizobiaceae). Relative abundance per sample was thereafter recalculated for major Hyphomicrobiales taxa. We evaluated the congruence in relative abundance estimations between *rpoB*- and 16S rRNA gene–based barcoding. We compared the relative abundance of major phyllosphere-associated Hyphomicrobiales genera in 41 leaf samples for which metabarcoding was performed with both 16S rRNA and *rpoB* genes [20]. To evaluate the congruence, we used a linear model on relative abundances after a Hellinger transformation to correct for rare taxa [14] and compared the observed estimate (slope) of the model to its expected distribution by realizing 10,000 random permutations of relative abundance values among leaf samples. Because relative abundances observed for a particular taxon could be site-dependent in this dataset [20], we constrained permutations within sites (SBL and MSH). We calculated p-values using the following formula: p=(b+1)/(m+1) where b was the number of expected values higher than the observed value and m, the number of permutations [32].

## RESULTS

### Phyllosphere-associated Hyphomicrobiales MAGs mostly belong to Methylobacteriaceae, Lichenihabitantaceae and two undescribed taxa

Phyllosphere-associated Hyphomicrobiales (syn. Rhizobiales) MAGs were previously identified in the phyllosphere of Canadian temperate forests (n=22; unpublished data; bioproject PRJEB93989), in mosses from the Canadian tundra (n=5; [18]), in lichens from a worldwide survey (n=43; [38]), in glacier sediments (n=1; [9]) and in mining soils (n=2; [9,24]; **Dataset 3**). We refined the taxonomic classification of phyllosphere-associated Hyphomicrobiales MAGs using a phylogenomic-based approach. Combining the 73 MAGs and 97 reference Hyphomicrobiales genomes covering the known phylogenomic diversity of phyllosphere-associated Hyphomicrobiales [13] (**Figure 1A; Figure S2; Dataset 3**), we identified 10,726 unique genes (hypothetical protein, repeat and mobile elements excluded). We considered 1,046 genes that were present in at least 80% of reference genomes (MAGs excluded) as candidate core genes. After excluding genes that were duplicated or missing data in more than 30% of assemblies (genomes and MAGs combined), we identified 824 core genes (37 rna and 787 protein-encoding genes; **Dataset 5**) for which we could retrieve the DNA sequence in at least 70% of genomes and MAGs, and thus infer a phylogenetic tree (RAxML, 1,000 replicated tree, GTRGAMMA model). Our consensus phyllosphere-associated Hyphomicrobiales coalescent tree combining the 824 core gene phylogenetic trees (with ASTRAL-III) was largely congruent with a previous phylogenomic-based classification of Hyphomicrobiales [13], except for the genus *Salinarimonas*, previously classified in Salinarimonadaceae, but embedded within the family Methylobacteriaceae according to our consensus tree. Our tree also provided a few additions for the genera *Lichenifustis*, *Methylocella* and *Hohaiirga*, confirming they belong to the families Lichenihabitantaceae, Beijerinckiaceae and Methylobacteriaceae, respectively. Our consensus tree supported the classification of most phyllosphere-associated Hyphomicrobiales MAGs into five monophyletic groups (branch support =1; **Figure 1B, Figure S2**). These groups corresponded to the families Methylobacteriaceae (n=18), Lichenihabitantaceae (n=27), Rhabdaerophilaceae (n=1) and to two undescribed taxa that we named according to the previous GTDB-based classification of phyllosphere-associated Hyphomicrobiales MAGs: namely: *RH-AL1* (n=21) and *JAJXWB01* (n=3). *RH-AL1* included only two reference genomes obtained from strains previously isolated from coal slag [45]. According to our consensus tree, it was a sister but distinct phylogenomic group from Beijerinckiaceae. *JAJXWB01* included no reference genome and was named according to a MAG identified in mine drainage sediments (unpublished data). According to our consensus tree, it was a sister but distinct phylogenomic group from Methylocystaceae and Rhodoblastaceae. No phyllosphere-associated Hyphomicrobiales MAG from related Hyphomicrobiales families could be identified (Alsobacteraceae, Boseaceae, Chelatococcaceae, Roseiarcaceae, Rhodoblastaceae, Methylocystaceae and Beijerinckiaceae). In Methylobacteriaceae, phyllosphere-associated Hyphomicrobiales MAGs from the genera *Enterovirga* (n=7) and *Methylobacterium* (n=11) were identified, but not from *Salinarimonas*, *Microvirga* or *Hohaiivirga*. In Lichenihabitantaceae, phyllosphere-associated Hyphomicrobiales MAGs were previously all assigned to Lichenihabitans according to the GTDB-based classification, but spanned all described genera according to our consensus tree, namely *Lichenihabitans* (n=8), *Lichenifustis* (n=6) and *Lichenibacterium* (n=13).

**Figure 1:**
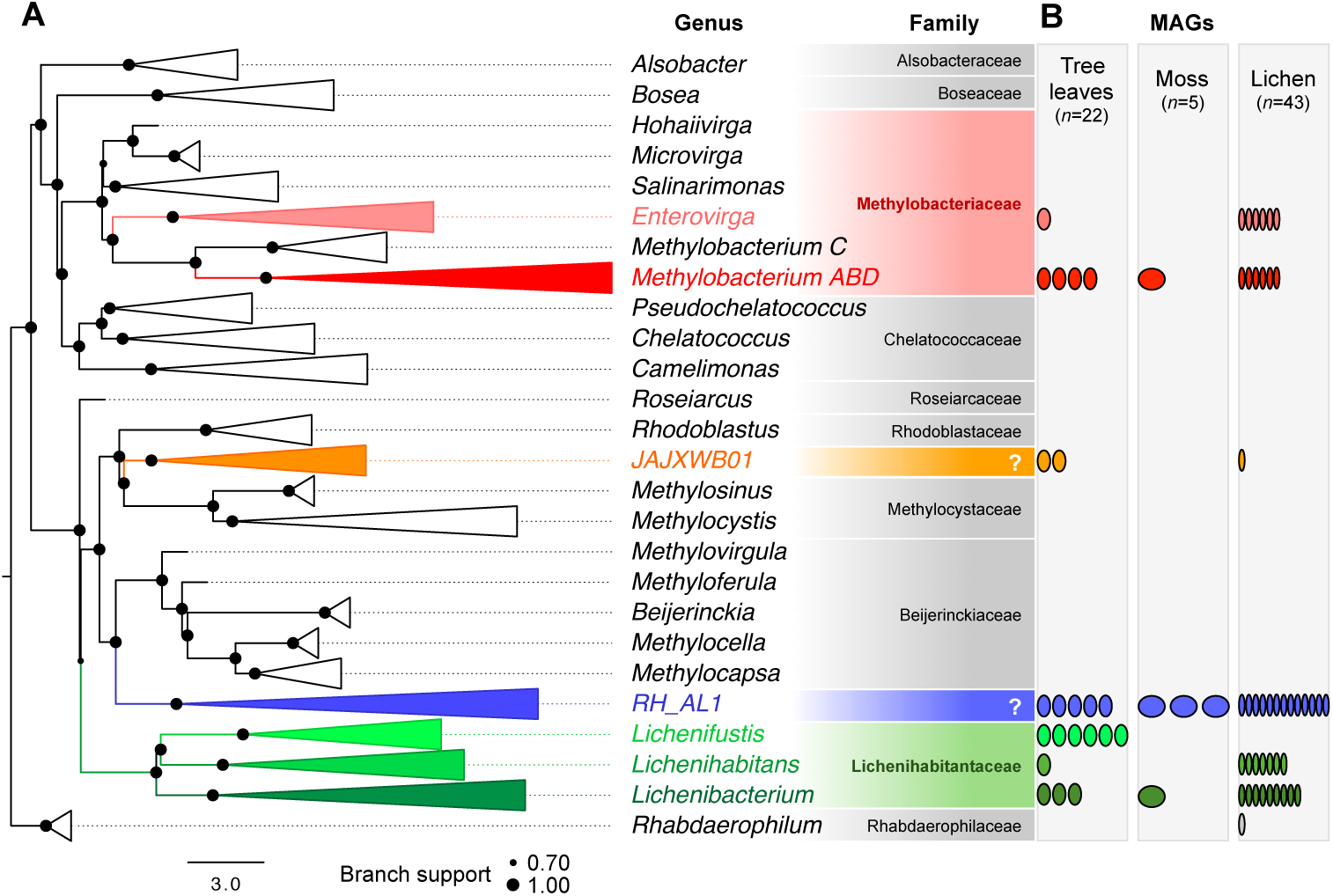
Phyllosphere-associated Hyphomicrobiales MAGs mostly belong to Methylobacteriaceae, Lichenihabitantaceae and to RH_AL1, an undescribed taxon related to Beijerinckiaceae. A) Phylogenomic tree of 96 reference genomes (NCBI ref_seq genomes) and 73 MAGs (tree leaves, moss and lichen). The consensus evolutionary tree was inferred from 824 core gene phylogenies combined with ASTRAL-III (See Figure S2 for detailed tree). The tree was rooted on *Rhabdaerophilum*. Scale represents coalescent time. Branch support represents local posterior probability. B) Distribution of tree leaf-, moss- and lichen-associated MAGs per Hyphomicrobiales genus. Each oval represents a MAG.

### Limits and improvements in the classification of phyllosphere-associated Hyphomicrobiales taxa with 16S rRNA metabarcoding and current databases

We verified the taxonomy of the phyllosphere-associated Hyphomicrobiales taxon *1174-901-12* in the previous SILVA database version 138.1, and identified as *Lichenibacterium* in version 138.2. A maximum-likelihood tree inferred from 24 complete 16S rRNA sequences used to define *1174-901-12* in the SILVA v138.1 database and from Lichenihabitantaceae type strains confirmed that this taxon corresponded to *Lichenibacterium* and to *Lichenifustis* (Figure S1; Dataset 2). In the tree, *1174-901-12* formed a monophyletic and strongly supported group with 16S rRNA genera *Lichenibacterium* and *Lichenifustis*, but not with their related counterpart *Lichenihabitans*.

We evaluated incongruences between taxonomic classifications of type strains from phyllosphere-associated Hyphomicrobiales with different databases: LPSN, NCBI, GTDB, and SILVA versions 138.1 and 138.2 (Table 1; Dataset 1). Rhizobiales, the illegitimate synonymous of Hyphomicrobiales, was still used in GTDB and in SILVA v138.1, but not in SILVA v138.2. In GTDB and in SILVA v138.1 and v138.2, most phyllosphere-associated Hyphomicrobiales families (Boseaceae, Methylobacteriaceae, Chelatococcaceae, Roseiarcaceae, Rhodoblastaceae, Methylocystaceae, Beijerinckiaceae and Lichenihabitantaceae) were incorrectly merged as Beijerinckiaceae, with the exception of two most basal genera *Alsobacter* and *Rhabdaerophillum*, which were identified as “Hyphomicrobiales *Incertae Sedis*” with SILVA. Genera *Pseudochelatococcus, Lichenibacterium* and *Rhodoblastus* were classified in LPSN as Beijerinckiaceae, Lichenibacteriaceae and Roseiarcaceae, respectively, while in NCBI, they were classified as Chelatococcaceae, Lichenihabitantaceae and Rhodoblastaceae, respectively (Table 1; Dataset 1). Genera *Lichenibacterium*, *Lichenihabitans* and *Lichenifustis* were merged as *Lichenihabitans* in GTDB and as *Lichenibacterium* in SILVA v138.2. *Methylorubrum* and *Methylobacterium* were considered as distinct genera in LPSN, NCBI and SILVA v138.2, but merged as *Methylobacterium* in GTDB and as *Methylobacterium-Methylorubrum* in SILVA v138.1.

We evaluated the power of the 16S rRNA gene as a marker to classify phyllosphere-associated Hyphomicrobiales taxa. We extracted 92 complete 16S rRNA sequences from reference phyllosphere-associated Hyphomicrobiales genomes (for which we validated the taxonomy using a phylogenomic approach; **Figure 1A**) that could be further used as a reference to refine phyllosphere-associated Hyphomicrobiales ASV taxonomy. Due to incomplete assemblies, no complete 16S rRNA gene sequence could be retrieved from most MAGs, but was recovered for JAJXWB01. A maximum-likelihood tree built from complete 16S rRNA gene sequences shows the limited power of this gene as a marker to infer the evolutionary history and taxonomy of phyllosphere-associated Hyphomicrobiales taxa (**Figure 2A**). In this tree, the families Methylobacteriaceae and Chelatococcaceae were not monophyletic, but deeply intermingled together. Within Beijerinckiaceae, the genera *Methylocystis* and *Methylocapsa* were not mophophyletic. According to the 16S rRNA gene tree, RH_AL1 and JAJXWB01 deeply branched within families Beijerinckiaceae and Methylocystaceae, respectively (**Figure 1A**), while these two undescribed groups formed sister but distinct clades from the two families in our phylogenomic tree (**Figure 2A**).

**Figure 2:**
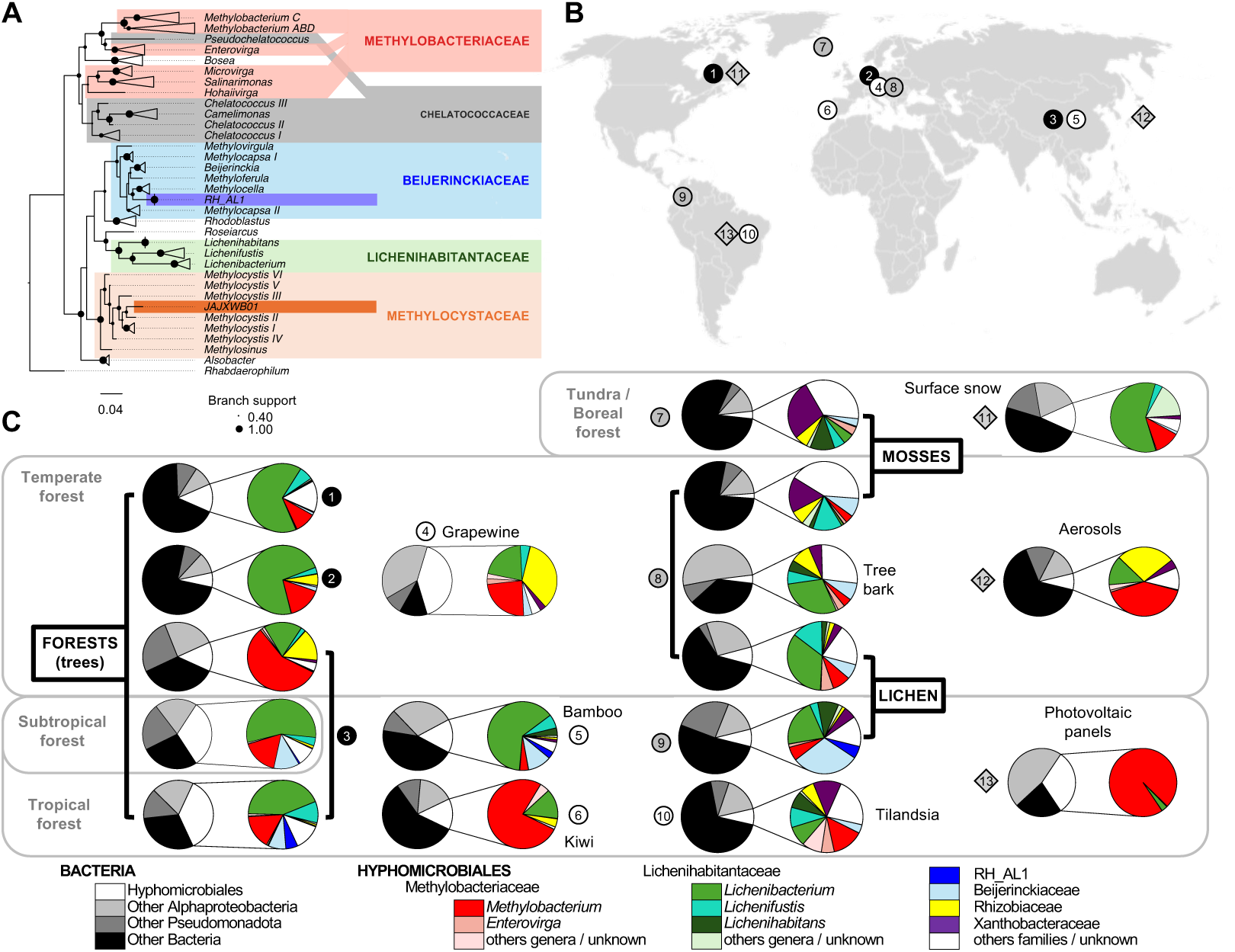
Phyllosphere is dominated worldwide by Methylobacteriaceae and Lichenihabitantaceae according to 16S rRNA gene metabarcoding. A) Maximum-Likelihood tree (1,000 bootstrap, GTRGAMMA model) from complete 16S rRNA gene sequences extracted from 96 phyllosphere-associated Hyphomicrobiales reference genomes (Figure 1). RH_AL1 is embedded within Beijerinckiaceae and JAJXWBA01 in Methylocystaceae. Methylobacteriaceae, Chelatococcaceae as well as some genera (*Chelatococcus*, *Methylocystis* and *Methylocapsa*) appear polyphyletic. B) World map showing the origin of BioProjects (numbers 1-13) presenting 16S rRNA metabarcoding data from the tree phyllosphere and potentially related microbial environments and sources (see **Dataset 4** for details). C) Bacterial abundance according to 16S rRNA barcoding, focusing on phyllosphere-associated Hyphomicrobiales families of the 13 BioProjects. Taxonomy from SILVA, refined with sequences from phyllosphere-associated Hyphomicrobiales reference genomes (see Fig. 1). Bioprojects were divided according to different climatic areas or substrates when applicable. For each category, abundances are shown for main taxonomic subdivisions other than Hyphomicrobiales (other Bacteria, Pseudomonadota and Alphaproteobacteria; left pie charts) and detailed for major families within Hyphomicrobiales and for all genera within Methylobacteriaceae and Lichenihabitantaceae (taxa with maximum relative abundance lower than 1% not showed, including Methylocystaceae and JAJXWBA01; right pie charts). For panels A and C, taxonomy follow phylogenomics classification (see Figure 1, Table 1).

We further explored the worldwide distribution and abundance of phyllosphere-associated Hyphomicrobiales taxa in the phyllosphere and related environments, using metabarcoding data from 13 studies based on the 16S rRNA gene (**Dataset 4; Figure 2B**). We identified 57,100 unique amplicon sequence variants (ASV) across 1998 samples. We first used SILVA 138.2 as a reference database to classify those ASVs at the order level. We identified 4,637 ASVs classified as Hyphomicrobiales, among which 3,089 were classified in phyllosphere-associated taxa, mostly Beijerinckiaceae (n=3,073). Because of SILVA databases limitations (**Table 1**), we used 92 complete 16S rRNA gene sequences extracted from complete phyllosphere-associated Hyphomicrobiales genomes (n=91) and MAG (n=1) as a database to reclassify the 3,089 ASVs at the family and genus level (**Dataset 6)**, according to the phylogenomic consensus tree (Figure 1A), with two exceptions. For family prediction, and according to the 16S rRNA gene phylogeny (Figure 2A), we classified JAJXWB01 within Methylocystaceae and RH_AL1 within Beijerinckiaceae (**Dataset 6**). Classification with SILVA, version 138.2 and with sequences extracted from reference genomes resulted in the same genus for 52.22% of ASVs (**Dataset 6**). Because most genera were classified within Beijerinckiaceae in SILVA database, we could not directly compare both methods for family classification. We thus deduced families in SILVA classification from the genus (e.g. by classifying *Methylobacterium* ASVs in Methylobacteriaceae instead of Beijerinckiaceae) and found that it matched family classification with sequences extracted from reference genomes for 81.57% of ASVs. Family could not be assigned with at least one method for 17.03% of ASVs. Among 1,338 ASVs identified as *Lichenibacterium* with SILVA, v 138.2 (43.31% of ASVs), we confirmed that 1,213 belonged to Lichenihabitantaceae using sequences from reference genomes as a database, and refined their classification in *Lichenibacterium* (701 ASVs), *Lichenifustis* (312), *Lichenihabitans* (121) and unknown Lichenihabitantaceae (79; **Dataset 6**).

### Lichenihabitantaceae and Methylobacteriaceae are dominant phyllosphere-associated Hyphomicrobiales taxa worldwide according to 16S rRNA metabarcoding

Worldwide, Pseudomonadota (formerly Proteobacteria) was the most abundant bacteria phylum in the phyllosphere of vascular, root-forming plants (excluding mosses and epiphytic plants), representing on average 56.1% of sequences identified as bacteria, or bacteria diversity [range: 25.1-87.1%] among Bioproject (**Dataset 7**; Figure 2C) and mostly consisted of Alphaproteobacteria (42.3 [16.7-78.7] % of Bacteria diversity). On average, Hyphomicrobiales dominated Alphaproteobacteria diversity in the phyllosphere (21.6 [6.8-40.6]% of total bacteria diversity) followed by Sphingomonadales (10.6 [2.2-21.0]%) and Acetobacterales (6.8[0.6-15.9%]). Hyphomicrobiales diversity in the phyllosphere was dominated by Lichenihabitantaceae (10.0 [2.0-19.7]% of total bacteria diversity) and by Methylobacteriaceae (5.8[0.7-11.8]% of total bacteria diversity). Other Hyphomicrobiales families, including Rhizobiaceae (2.2 [0.1-14.0]% of total bacteria abundance) and Beijerinckiaceae+RH_AL1 (1.7 [0.0-5.0]% of total bacteria diversity), could be occasionally detected at the surface of tree leaves, but barely Methylocystaceae+JAJXWB01 (0.3 [0.0-0.9]% of total bacteria diversity). In the phyllosphere, ASVs identified as Methylobacteriaceae mostly corresponded to *Methylobacterium* (5.3 [0.6-11.0]% of total bacteria diversity) and ASVs identified as Lichenihabitantaceae mostly corresponded to *Lichenibacterium* (8.7 [2.0-17.5]% of total bacteria diversity) and *Lichenifustis* (1.1[0.0-3.7]%). ASVs classified as RH_AL1 were barely observed (0.4[0.0-2.0]% of total bacteria diversity) and JAJXWB01 was almost never detected (< 0.1%). On average, Hyphomicrobiales represented a smaller proportion of bacterial diversity in lichen habitats (8.1[7.8-8.4]% of total bacteria diversity) and moss habitats (2.9[2.4-3.5]% of total bacteria diversity) compared to the phyllosphere of vascular plants (21.6 [6.8-40.6]% of total bacteria diversity). In lichen, the Hyphomicrobiales were still dominated by Lichenihabitantaceae lichen (3.6[2.8-4.4]% of total bacteria diversity). Notable proportions of Lichenihabitantaceae were also detected in snow surface (10.6% of total bacteria abundance), the surface of tree bark (1.6%), in epiphytic plants (1.9%), and in aerosols (1.0%). Methylobacteriaceae was one of the dominant bacterial families detected at the surface of photovoltaic panels (30.2% of total bacteria abundance) and was also detected in aerosols (3.4%), on epiphyte plants (2.1%) and on snow surface (1.6%).

### Comparison of 16S rRNA and *rpoB*–based barcoding for the identification of phyllosphere-associated Hyphomicrobiales taxa

We explored the potential of the *rpoB* gene, which we previously adapted to metabarcoding to evaluate the diversity of *Methylobacterium* in the phyllosphere, as a broader taxonomic marker for Hyphomicrobiales [20]. We extracted 102 complete *rpoB* sequences from reference phyllosphere-associated Hyphomicrobiales genomes (n=95) and MAGs (n=7), which could be further used as a reference to refine the taxonomy of phyllosphere-associated Hyphomicrobiales ASVs. A maximum-likelihood phylogenetic tree built from complete *rpoB* gene nucleotide sequences shows that this gene is a reliable marker to infer the evolutionary history and taxonomy of phyllosphere-associated Hyphomicrobiales taxa (**Figure S2**). In this tree, all families predicted by the phylogenomic-based consensus tree (**Figure 1A)** were monophyletic (branch support > 0.90). Within phyllosphere-associated Hyphomicrobiales families (Lichenihabitantaceae, Methylobacteriaceae), all genera predicted by the phylogenomic-based consensus tree (**Figure 1A)** were monophyletic (branch support > 0.81).

We evaluated the abundance of Hyphomicrobiales taxa from *rpoB* metabarcoding previously developed to target Methylobacteriaceae and performed on 184 leaf samples from two temperate forests in Quebec [20]. ASVs were mostly assigned to Hyphomicrobiales (average per sample; 95.6%; range per forest: 93.8-97.7%) and Caulobacterales (3.9[1.7-5.8]%; Dataset 8). We refined previous taxonomic assignments only for Hyphomicrobiales ASVs, using *rpoB* sequences extracted from reference genomes and MAGs as a reference. In Hyphomicrobiales, ASVs were mostly classified in three families: Methylobacteriaceae (24.6[12.2-39.0%]) - dominated by *Methylobacterium* (23.4[11.1-37.7]%), RH_AL1 (19.8[13.1-25.5]%) and Lichenihabitantaceae (36.7[31.1-41.5]%) - dominated by *Lichenibacterium* (31.9[23.0-39.6]%) and *Lichenifustis* (4.3[1.1-8.0]%). *Enterovirga* (Methylobacteriaceae) and JAJXWB01 could only be detected at very low abundances (0.2[0.1-0.3]% and 0.1[0.0-0.3]%, respectively) (Figure S2B). We compared the relative abundance of Phyllosphere-associated Hyphomicrobiales genera in 41 leaf samples for which metabarcoding was performed with both 16S rRNA and *rpoB* genes, namely *Methylobacterium*, *Enterovirga*, *Lichenifustis*, *Lichenibacterium,* RH_AL1 and JAJXWB01 (**Figure S3C)**. For these taxa, we observed a significant correlation between relative abundances estimated by 16S rRNA gene and *rpoB* gene barcoding (R^2^= 0.529, p<0.001; **Figure S3C).** This correlation was not significantly different than expected after 10,000 permutations of relative abundances among leaf samples, within or among sites (*p* > 0.05; **Figure S3D)**, indicating that it was mostly explained by the contrast between highly frequent and rare taxa (*Lichenibacterium* and *Methylobacterium* vs *Lichenifustis* and *Enterovirga*), consistently observed among leaf samples, and by differences between forests (SBL vs MSH).

## DISCUSSION

To clarify the distribution and composition of phyllosphere-associated Hyphomicrobiales, we examined over 1,900 phyllosphere 16S rRNA gene datasets from 13 studies spanning multiple continents. This meta-analysis confirmed that *Hyphomicrobiales* constitute a significant fraction of phyllosphere bacterial communities, with the families *Methylobacteriaceae* and *Lichenihabitantaceae* being dominant across environments (Figure 2C). Notably, these groups also co-occurred in related aerial habitats such as lichen and moss surfaces, snow, and aerosols, suggesting they may form part of a larger aeroterrestrial biome. The identification of Lichenihabitantaceae, which was solely isolated and characterized in lichen [30], as a dominant taxon of the phyllosphere of trees, has profound implications for understanding microbial colonization dynamics in both the phyllosphere and extreme environments. The ecological distribution of Lichenihabitantaceae on exposed surfaces, leaves and potentially in snow, suggests its resilience to environmental stresses, marking it as a pioneer organism capable of colonizing newly exposed surfaces. This resilience aligns with potential pioneering capacities of *Lichenibacterium*, enabling it to establish communities on surfaces exposed to UV and nutrient scarcity [30]. For instance, metagenomics from lichens confirmed the abundance of *Lichenibacterium* and suggested that these bacteria may play a key role in providing essential nutrients to their fungal symbiotic partners in harsh environments [1]. Furthermore, *Lichenibacterium*’s dominance as an early leaf colonizer implies that it could facilitate subsequent microbial community development, potentially influencing plant health and ecosystem functioning.

Our analyses also highlighted the limitations of the 16S rRNA gene for resolving phyllosphere-associated Hyphomicrobiales diversity. Key families such as *Methylobacteriaceae* appeared polyphyletic based on 16S sequences, and clade-level patterns were often ambiguous or inconsistent across classification schemes (Figure 2A), illustrating the low resolution of the 16S rRNA gene in inferring taxonomy and evolutionary history in some bacterial taxa [2,7,15]. We also confirmed that Lichenihabitantaceae, especially the genera *Lichenifustis* and *Lichenibacterium*, corresponded to *1174-901-12* (Figure S1; Table 1), a undescribed Hyphomicrobiales taxon identified in previous but widely used SILVA version (v 138.1), and frequently associated with the phyllosphere [3,16,20,23,36,44,46]. The ecological and functional explanation of *1174-901-12* high abundance on exposed surfaces, especially plant leaves, remained largely elusive. Its identification as Lichenihabitantaceae paths the way to a re-examination of previous environmental studies based on 16S rRNA metabarcoding and to a better understanding of its ecological role.

By comparing main taxonomic databases used in microbial ecology for the classification of phyllosphere-associated Hyphomicrobiales, we highlighted another major source of biases and potential misinterpretation of the diversity and ecological role of this group (Table 1). A striking example concerns the merging of distinct families such as *Boseaceae*, *Methylobacteriaceae*, *Chelatococcaceae*, *Roseiarcaceae*, *Rhodoblastaceae*, *Methylocystaceae*, *Beijerinckiaceae*, and *Lichenihabitantaceae* under a single family (*Beijerinckiaceae*) in GTDB and SILVA. Such simplifications mask the phylogenetic and ecological distinctiveness of these lineages and obscure evolutionary signals relevant to host association and adaptation within the phyllosphere. Inconsistencies were especially marked for dominant phyllosphere-associated families Lichenihabitantaceae and Methylobacteriaceae, illustrating the inertia of nomenclatural changes across widely used databases. In GTDB and SILVA v138.2, *Lichenibacterium*, *Lichenihabitans*, and *Lichenifustis* were merged under a single genus, collapsing taxonomic distinctions supported by genomic data (Figure 1), while these genera were classified in distinct families in LPSN. The inconsistent treatment of *Methylobacterium* and *Methylorubrum* across databases further illustrates the consequences of divergent interpretations of recent taxonomic revisions [2]. Such inconsistencies may lead to underestimation of genus-specific ecological adaptations to the phyllosphere and reduce the resolution of community analyses based on amplicon or metagenomic data.

To overcome these limitations, we applied a genome-resolved phylogenomic approach using core single-copy genes to analyze a representative set of 73 phyllosphere-associated Hyphomicrobiales MAGs alongside 97 reference *Hyphomicrobiales* genomes (Figure 1, Figure S2). This approach provided a clearer taxonomic framework, revealing that the majority of phyllosphere-associated Hyphomicrobiales MAGs clustered into four robust clades: *Methylobacteriaceae*, *Lichenihabitantaceae*, and two previously undescribed but well-supported lineages—RH-AL1 and JAJXWB01. RH-AL1 is represented by two reference isolates from coal slag and is related to *Beijerinckiaceae* [45], but forms a distinct and phylogenetically diverse lineage, including MAGs from the phyllosphere, lichens and from mosses. JAJXWB01 appears completely uncultured and forms a related but distinct lineage from *Methylocystaceae*, including MAGs from lichen and from the phyllosphere. Interestingly, RH-AL1 reference isolates, like many members of the Beijerinckiaceae and Methylocystaceae families, are methylotrophic bacteria [45], a trait shared with *Methylobacterium* and allowing leaf-associated bacteria to use methanol released by plant as carbon source [39]. However, no MAG was placed within Methylocystaceae or Beijerinckiaceae, suggesting a relatively narrow and structured set of phyllosphere-associated Hyphomicrobiales lineages adapted to phyllosphere environments. A similar narrow ecological distribution was observed for Methylobacteriaceae, with phyllosphere-associated Hyphomicrobiales MAGs only identified within *Enterovirga* and *Methylobacterium*, but not in other genera (Figure 1). *Enterovirga* was traditionally associated with animal feces or aerosols [11,22], but its detection in the phyllosphere and in lichens suggest a larger ecological niche. In *Methylobacterium*, MAGs were only detected in traditionally leaf-associated phylogroups A, B and D, but not in the most ancestral group C, confirming secondary leaf-association during the evolution of this genus [2,21]. The occurrence of phyllosphere-associated traits should be further investigated in phyllosphere-associated Hyphomicrobiales genera, for instance by screening the occurrence of methylotrophic pathways in phyllosphere-associated Hyphomicrobiales genomes and MAGs. The ability for these bacteria to effectively metabolize one-carbon compounds like methanol should be next validated by culturomics and phenotypic screening, in particular within taxa traditionally not associated with the phyllosphere like Lichenihabitantaceae and *Enterovirga*, or poorly represented in genomic databases and strain collections like JAJXWB01.

To complement the phylogenomic analysis of phyllosphere-associated Hyphomicrobiales, we evaluated the utility of the alternative marker gene *rpo*B that we previously used to improve resolution in *Methylobacterium* diversity characterization in the phyllosphere [20]. Unlike the 16S rRNA gene, *rpoB*-based phylogenies recovered phyllosphere-associated Hyphomicrobiales family- and genus-level relationships with much greater accuracy and congruence with genome-based placements (Figure S3). In particular, using *rpoB* amplicon sequencing revealed the strong prevalence of *Methylobacterium* and *Lichenibacterium*, supporting their dominance inferred from metagenomic data. Furthermore, the RH-AL1 clade, which was poorly detected by 16S rRNA barcoding, accounted for a large fraction of *Hyphomicrobiales* reads in *rpoB*-based profiles, of roughly the same abundance as *Methylobacterium* and *Lichenibacterium*, highlighting the *rpoB* marker’s superior sensitivity and specificity for uncovering cryptic diversity in phyllosphere-associated Hyphomicrobiales. We cannot rule out that the observed relative abundances are biased, as the *rpoB* marker was optimized for *Methylobacteriaceae*; however, the opportunity to design a more generalist marker is limited by the high polymorphism of this gene, and metagenomics ultimately provides a more comprehensive alternative.

Our study provides a revised and integrative view of phyllosphere-associated *Hyphomicrobiales*, revealing both the global dominance of known families such as *Methylobacteriaceae* and *Lichenihabitantaceae*, and the existence of undescribed but recurrent clades. We show that relying solely on the 16S rRNA gene obscures key phylogenetic and ecological patterns of bacterial communities in the phyllosphere, particularly due to its limited resolution and lack of congruences across reference databases. By integrating *rpoB* markers and genome-resolved phylogenetics, we propose a more robust framework to track phyllosphere-associated Hyphomicrobiales diversity and guide future ecological and evolutionary studies. Looking forward, the characterization of the novel RH-AL1 and JAJXWB01 lineages, as well as a better delineation of the genera within *Lichenihabitantaceae*, are important research priorities. Combining long-read metagenomics, transcriptomics, and targeted functional assays will further allow us to link genome evolution to ecological adaptation, helping to uncover the roles these widespread Hyphomicrobiales play in the phyllosphere and beyond.

## Funding information

JBL and SWK were supported by NSERC Discovery Grants RGPIN-2024-05825 and RGPIN-2025-05608 respectively.

## Acknowledgments

NIngeborg Klarenberg and Wisnu Adi Wicaksono for providing Metadata; Marta Alonso Garcia and Zihui Wang for discussion.

## Data availability

Code and data are publicly available on Github repository: JBLED/Hyphomicrobiales-phyllosphere

## Notes

https://github.com/JBLED/Hyphomicrobiales-phyllosphere

### Competing Interest Statement

The authors have declared no competing interest.

### Summary of Updates

Major Changes in This Version - Rewrote the introduction and abstract to clarify objectives and the impact of 16S rRNA-based biases. - Added a new comparative analysis of taxonomic inconsistencies across LPSN, GTDB, NCBI, and SILVA, with corresponding additions to Results and Discussion (Table 1) - Updated all taxonomic assignments, figures, tables, and supplementary materials to SILVA v138.2. - Standardized nomenclature across the manuscript. - Clarified the interpretation of rpoB amplicon data, including potential primer bias and marker limitations. - Implemented all additional minor revisions for clarity, consistency, and formatting.

