## Supplementary material for "A phylogenomic and metagenomic meta-analysis of bacterial diversity in the phyllosphere lifts a veil on Hyphomicrobiales dark matter": Figure S1

**Figure S1: *Lichenibacterium* corresponds to *1174-901-12*.** A maximum-likelihood tree (1,000 bootstraps, GTRGAMMA model) built from complete 16S rRNA gene nucleotide sequences (1379 bp) shows that the genus annotated as *1174-901-12* in the SILVA Database (v138.1) corresponds to lichen-associated Hyphomicrobiales genera *Lichenibacterium* and *Lichenifustis*. NJ tree (1,000 bootstrapps) rooted on *Roseiarcus.* Nodes supported by less than 60% of bootstraps were collapsed.

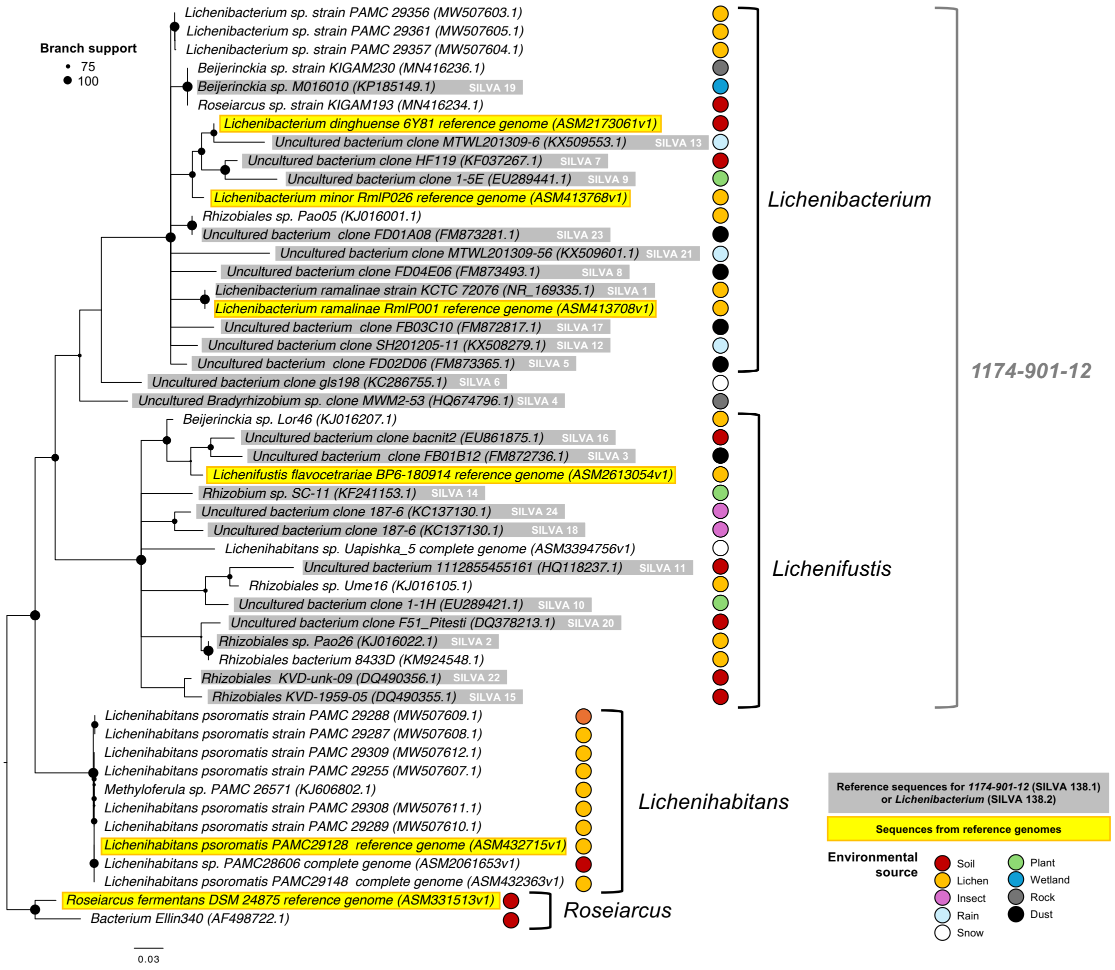
