## Supplementary material for "A phylogenomic and metagenomic meta-analysis of bacterial diversity in the phyllosphere lifts a veil on Hyphomicrobiales dark matter": Figure S2

**Figure S2: Detailed phylogeny of Phyllosphere-associated Hyphomicrobiales genomes and MAGs.** Phylogenomic tree of 98 reference genomes (NCBI ref_seq genomes) and 72 MAGs (tree leaves, moss and lichen). The consensus evolutionary tree was inferred from 824 core gene phylogenies combined with ASTRAL-III. The tree was rooted on *Rhabdaerophilum*. Scale represents coalescent time. Branch support represents local posterior probability.

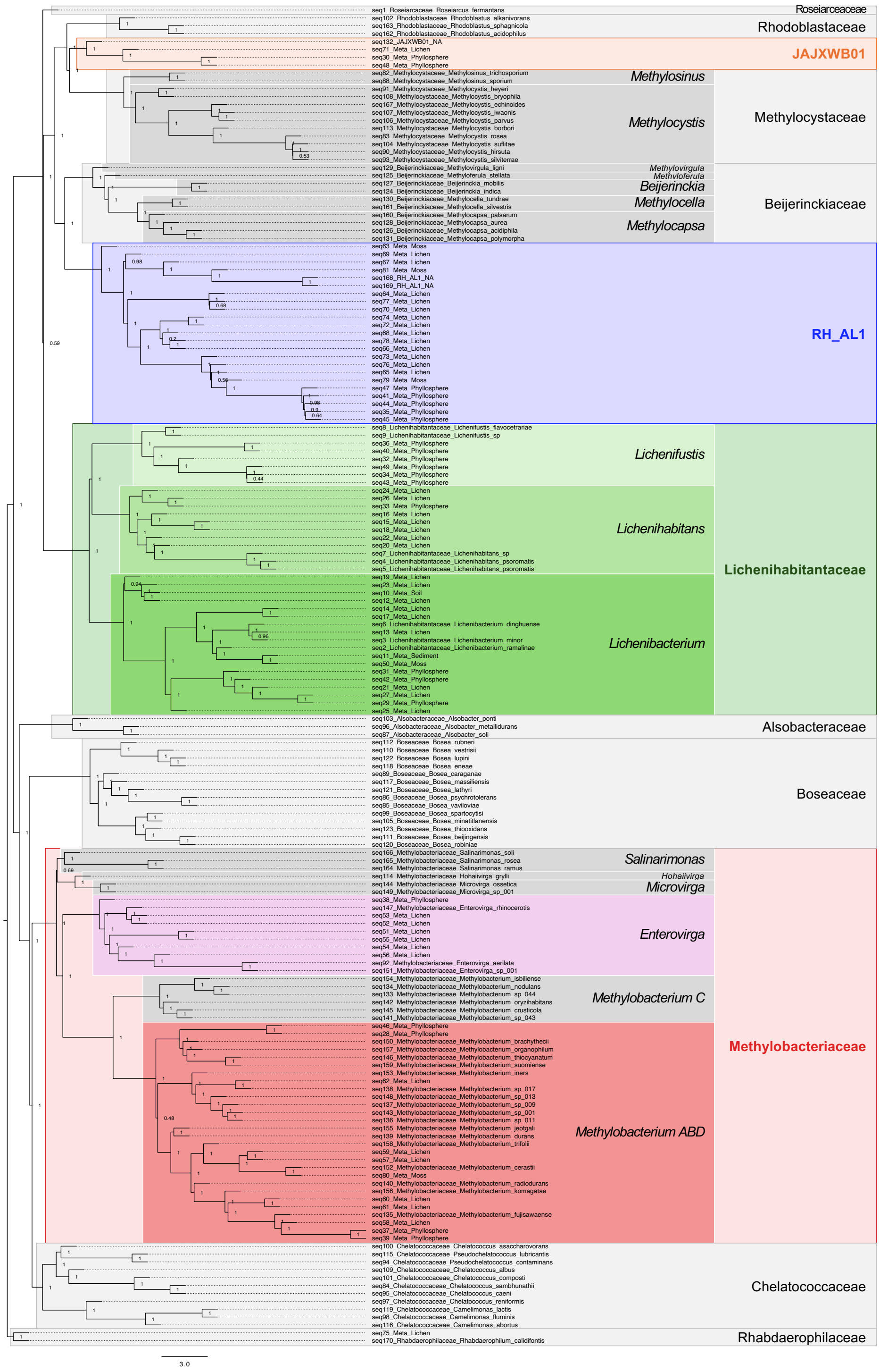
