## Supplementary material for "A phylogenomic and metagenomic meta-analysis of bacterial diversity in the phyllosphere lifts a veil on Hyphomicrobiales dark matter": Figure S3

**Figure S3: Comparison of barcoding from 16S rRNA and *rpoB* genes, data from Leducq et al. 2022 [20].** A) Maximum-Likelihood tree (1,000 bootstrap) from complete *rpoB* nucleotide sequences extracted from 102 phyllosphere-associated Hyphomicrobiales genomes and MAGs. Branches assigned to the same genera were collapsed. B) Relative abundance estimated from *rpoB* barcoding of *Lichenibacterium*, *Lichenifustis*, *Enterovirga*, *Methylobacterium*, RH-AL1 and JAJXWB01 in 184 tree leaf samples from two Canadian temperate forests (Mont St-Hilaire - MSH and Station Biologique des Laurentides - SBL). Bars indicate means for each genus (color codes from tree in panel A). C) Comparison of relative abundances of four phyllosphere-associated Hyphomicrobiales genera that could be identified with both markers in 41 leaf samples from MSH and SBL. Each dot represents the relative abundances (Hellinger transformation) of a taxon in each leaf sample estimated by *rpoB* barcoding (Y-axis) and *16S rRNA* barcoding (X-axis; log scale). The linear regression between both method (red line) and distribution of this regression after 10,0000 permutations (grey; see panel D) are represented. D) Distribution of correlation coefficients in the comparison between relative abundances estimated from 16S rRNA gene and *rpoB* gene barcoding, after 10,000 random permutations of relative abundances among leaf samples, among and within sites (light grey) or only within sites (dark grey).

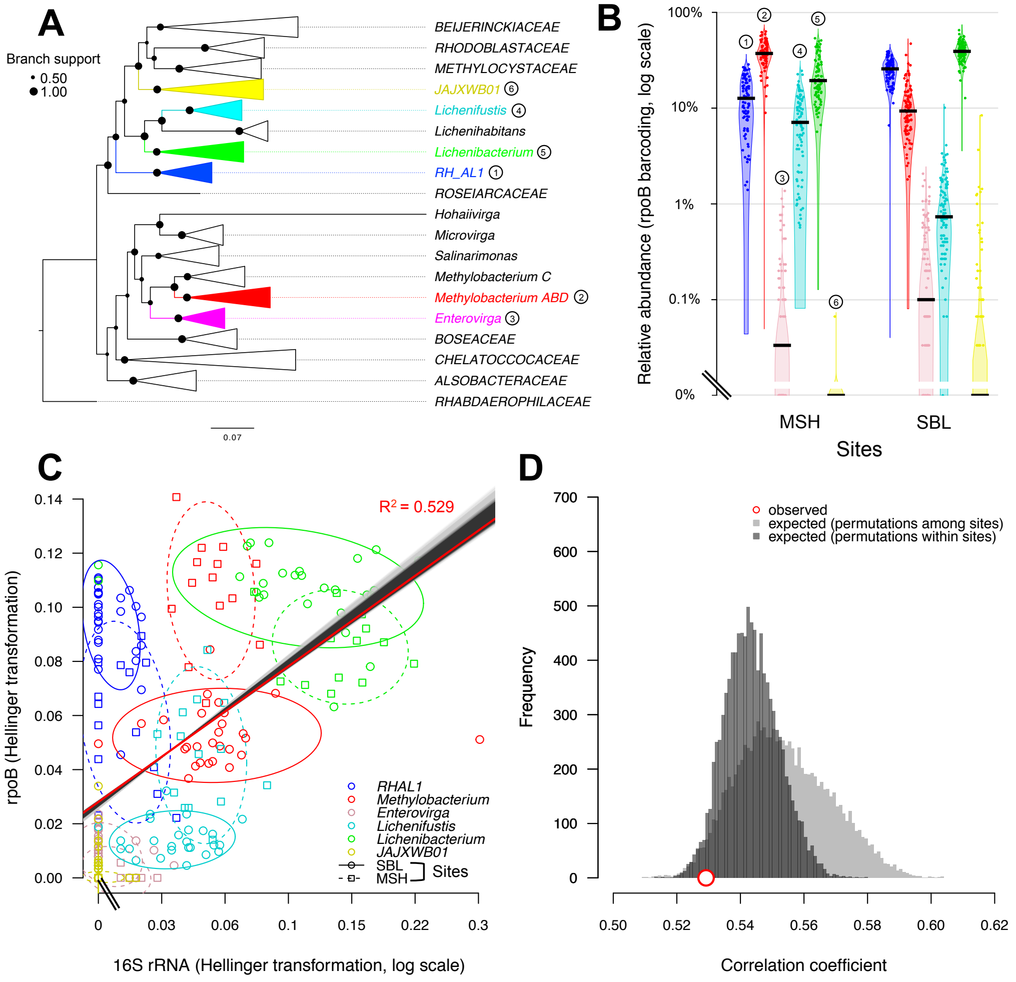
